# Parvalbumin interneuron dysfunction in a thalamus - prefrontal cortex circuit in Disc1 deficiency mice

**DOI:** 10.1101/054759

**Authors:** Kristen Delevich, Hanna Jaaro-Peled, Mario Penzo, Akira Sawa, Bo Li

## Abstract

Two of the most consistent findings across disrupted-in-schizophrenia-1 (DISC1) mouse models are impaired working memory and reduced number or function of parvalbumin interneurons within the prefrontal cortex. While these findings suggest parvalbumin interneuron dysfunction in DISC1-related pathophysiology, to date, cortical inhibitory circuit function has not been investigated in depth in DISC1 deficiency mouse models. Here we assessed the function of a feedforward circuit between the mediodorsal thalamus (MD) and the medial prefrontal cortex (mPFC) in mice harboring a deletion in one allele of the Disc1 gene. We found that the inhibitory drive onto layer 3 pyramidal neurons in the mPFC was significantly reduced in the Disc1 deficient mice. This reduced inhibition was accompanied by decreased GABA release from local parvalbumin, but not somatostatin, inhibitory interneurons, and by impaired feedforward inhibition in the MD-mPFC circuit. Our results reveal a cellular mechanism by which deficiency in DISC1 causes neural circuit dysfunction frequently implicated in psychiatric disorders.

---

Mounting evidence suggests that dysregulation of excitation-inhibition (E-I) balance in neural circuits may be a pathophysiological feature of schizophrenia (Lewis et al., 2011; Lisman, 2012). Data from rodent studies support the role of prefrontal fast-spiking parvalbumin (PV) inhibitory interneurons (INS) in cognitive tasks that are impaired in schizophrenia, particularly working memory (Cho et al., 2015; Murray et al., 2015). Cognitive impairment is commonly seen in first-degree relatives of individuals with schizophrenia (Cannon et al., 2000; Myles-Worsley and Park, 2002; Snitz et al., 2006), suggesting that cognitive functions such as working memory, and the brain structures and circuits that subserve them, are highly heritable. Disrupted-in-schizophrenia-1 (DISC1) has emerged as a rare, but penetrant genetic risk factor for psychopathology (Millar et al., 2000), and DISC1 polymorphisms have been reported to correlate with measures of cognitive performance and frontal lobe structure (Cannon et al., 2005; Carless et al., 2011; Hennah et al., 2005; Palo et al., 2007).Work in mouse models has revealed the importance of the Disc1 gene in neurodevelopment (Kamiya et al., 2005; Mao et al., 2009; Niwa et al., 2010), synaptic function (Hayashi-Takagi et al., 2010; Maher and LoTurco, 2012; Sauer et al., 2015; Seshadri et al., 2015; Wang et al., 2011; Wei et al., 2015), and cognitive processing (Brandon and Sawa, 2011). Notably, working memory impairments are consistently reported across DISC1 mouse models tested (Brandon and Sawa, 2011; Clapcote et al., 2007; Koike et al., 2006; Kvajo et al., 2008; Lee et al., 2013; Li et al., 2007; Lipina et al., 2010; Niwa et al., 2010). Furthermore, a variety of DISC1 mouse models also exhibit reduced prefrontal PV expression (Ayhan et al., 2011; Hikida et al., 2007; Ibi et al., 2010; Lee et al., 2013; Niwa et al., 2010; Shen et al., 2008), suggesting that PV INs may be particularly affected by DISC1 perturbation. To improve our understanding of the role of DISC1 in prefrontal cortical circuit function, we examined local and long-range circuits of the mPFC in mice heterozygous for a deficiency (df) allele of the DISC1 gene (DISC1df/+) (Seshadri et al., 2015; Shahani et al., 2015). We found that DISC1df/+ mice exhibited a specific deficit in inhibitory synaptic transmission. This deficit is expressed as reduced frequency of miniature inhibitory post-synaptic currents (mIPSCs) onto layer 3 (L3) pyramidal neurons (PNs), increased paired-pulse ratio of IPSCS onto these PNs driven by local PV INS, and impaired feedforward inhibition in the mediodorsal thalamus-mPFC circuit that is mediated by PV INS in the mPFC (Delevich et al., 2015). These data support the hypothesis that the genetic risk factor DISC, impacts the function of PV INS, and provide insight into the synaptic and circuit mechanisms of DISC1-related cognitive dysfunction.

## I. METHODS

### i. Animals

Mice were group housed under a 12-h light-dark cycle (9 a.m. to 9 p.m. light), with food and water freely available. Both male and female mice were used. All procedures involving animals were approved by the Institute Animal Care and Use Committees of Cold Spring Harbor Laboratory and conducted in accordance to the US National Institute of Health guidelines. The PV-Cre (jaxmice.jax.org/strain/008069.html), SOM-Cre (jaxmice.jax.org/strain/013044.html), and Ail4 (www.jax.org/strain/007914) mice were described previously (Hippenmeyer et al., 2005; Madisen et al., 2010; Taniguchi et al., 2011). We recently generated the Disc1 deficiency mice, which harbor a deletion (6.9 kb) encompassing the first 3 exons of the Disc1 gene (Seshadri et al., 2015). The majority of DISC1 isoforms are abolished in mice homozygous for the Disc1 deficiency (df) allele (Disc1df/df) (Seshadri et al., 2015). We used heterozygous deficiency of Discl in mice (Disc1df/+) to model DISC1 genetic deficits that could be seen in humans. All mice have been bred onto C57BL/6N background for at least 5 generations.

### ii. Viral vectors

Adeno-associated virus (AAV) vectors AAV-CAG-ChR2(H134R)-YFP and AAV-eF1a-DIO-ChR2(H134R)-YFP were produced as AAV2/9 serotype by the University of North Carolina Vector Core (Chapel Hill, NC) and has been previously described (Delevich et al., 2015; Zhang et al., 2007). All viral vectors were stored in aliquots at −80° C until use.

### iii. Stereotaxic surgery

Mice aged postnatal day 40 to 45 (P40-P45) were used for all surgeries. Unilateral viral injections were performed using previously described procedures (Li et al., 2013) at the following stereotaxic coordinates: MD,-1.58 mm from Bregma, 0.44 mm lateral from midline, and 3.20 mm vertical from cortical surface; dorsal mPFC: 1.94 mm from Bregma, 0.34 mm lateral from midline, and 0.70 mm vertical from cortical surface. Surgical procedures were standardized to minimize the variability of AAV injections. To ensure minimal leak into surrounding brain areas, injection pipettes remained in the brain for 5 min post-injection before being slowly withdrawn. The final volume for AAV-CAG-ChR2(H134R)-YFP injected into MD was 0.3-0.35 μL, and for AAV-eFla-DI0-ChR2(H134R)-YFP injected into dorsal mPFC was 0.5 μL. The titer for the viruses was 1012 viral particles/mL. Viruses were allowed to incubate for 2-4 weeks for optimal expression.

### iv. Electrophysiology

Some of the control data from WT mice used for comparing with Disc1df/+ mice(appearing in Fig. 4,5 and 6) were previously reported in Fig. 12, and 4 of (Delevich et al., 2015).

Mice were anaesthetized with isoflurane and decapitated, whereupon brains were quickly removed and immersed in ice-cold dissection buffer (ll0.0mM choline chloride, 25.0 mm NaHC03,1.25 mm NaH2 P04, 2.5 mm KCl, 0.5 mm CaCl2, 7.0 mm MgCl2, 25.0 mm glucose,ll.6mM ascorbic acid and 3.1 mm pyruvic acid, gassed with 95% 02 and 5% C02). Coronal slices (300 μm in thickness) containing mPFC were cut in dissection buffer using a HM650 Vibrating Microtome (Thermo Fisher Scientific), and were subsequently transferred to a chamber containing artificial cerebrospinal fluid (ACSF) (ll8mM NaCl, 2.5 mm KCl, 26.2 mm NaHC03, lmM NaH2 P04, 20 mm glucose, 2mM MgCl2 and 2mM CaCl2, at 34°C, pH 7.4, gassed with 95% 02 and 5% C02). After ~30 min recovery time, slices were transferred to room temperature and were constantly perfused with ACSF.

The internal solution for voltage-clamp experiments contained l40mM mm potassium gluconate, l0mM HEPES, 2mM MgCl2, 0.5mM CaCl2, 4mM MgATP, 0.4 mm Na3GTP, l0mM Na2-Phosphocreatine, l0mM BAPTA, and 6 mm QX-314 (pH 7.25, 290 m0sm). Elec-trophysiological data were acquired using pCLAMP 10 software (Molecular Devices). mIPSCs were recorded in the presence of tetrodotoxin l μm, APV 100 μm, and CNQX 5μM. mEPSCs were recorded in the presence of tetrodotoxin l μm and picrotoxin 100 μm. Data were analyzed using Mini Analysis Program (Synaptosoft). For the mIPSCs and mEPSCs, we analyzed the first 300 and 250 events, respectively, for each neuron. The parameters for detecting mini events were kept consistent across neurons, and data were quantified blindly with regard to the genotypes.

To evoke synaptic transmission by activating ChR2, we used a single-wavelength LED system (λ = 470 nm, CoolLED.com) connected to the epifluorescence port of the 0lympus BX5l microscope. To restrict the size of the light beam for focal stimulation, a built-in shutter along the light path in the BX51 microscope was used. Light pulses of 0.5-1 ms triggered by a TTL (transistor-transistor logic) signal from the Clampex software (Molecular Devices) were used to evoke synaptic transmission. The light intensity at the sample was ~ 0.8 mW/mm^2^. Electrophysiological data were acquired and analyzed using pCLAMP 10 software (Molecular Devices). IPSCS were recorded at 0 mV holding potential in the presence of 5 μm CNQX and 100 μm AP-5. Light pulses were delivered once every 10 seconds, and a minimum of 30 trials were collected. In paired-pulse recordings, 2 light pulses separated by 50, 100, or 150 ms were delivered. In cases that the first IPSC did not fully decay to baseline before the onset of the second IPSC, the baseline of the second IPSC was corrected before the peak was measured. To measure the kinetics of the IPSCS, averaged sweeps collected at the 150 ms interval were normalized, and the decay time constant and half-width were measured using automated procedures in the AxoGraph X 1.5.4 software.

To determine IPSC reversal potential (EIPSC), IPSCS were recorded at varying holding potentials (20 mV steps) in the presence of 5 μm CNQX and 100 μm APV to block AMPA receptors and NMDA receptors, respectively. IPSC amplitude was measured, and a linear regression was used to calculate the best-fit line, and the x-intercept was used as the EIPSC. Under our recording conditions, the EIPSC was ~ -60 mV. Therefore, in the excitation/inhibition ratio (E/I) experiments, we recorded EPSCS at -60 mV and IPSCS at 0 mV holding potential. The only drug used for the E/I experiments was APV l00μM. In these experiments we used the same light intensity for evoking both IPSCS and EPSCS. In addition, we used similar stimulation regime for WT and Disc1df/+ mice, such that the peak amplitudes of IPSCS are comparable between genotypes.

For the experiments in which we optogenetically stimulated the MD axons in the mPFC, mice were excluded if the extent of infection in the MD was too large and leaked into surrounding brain regions. Rodent MD lacks interneurons; therefore all ChR2 infected neurons are expected to be relay projection neurons (Kuroda et al., 1998).

The latency and 10–90% rise-time of EPSCS and IPSCS were calculated from either the averaged trace or individual sweeps for each cell using automated procedures in the AxoGraph X 1.5.4 software. ESPC and IPSC onset latency was calculated as the time from stimulation onset to 10% rise time, with EPSC-IPSC delay calculated as the difference. The 10% rise time has been reported to be a more reliable measure of delay to onset, as it minimizes the contribution of EPSC and IPSC rise time differences that are reflected in the time to peak (Mittmann et al., 2005).

### v. Data analysis and statistics

All statistical tests were performed using 0rigin 9.0 (0rigin-Lab, Northampton, MA) or GraphPad Prism 6.0 (GraphPad Software, La Jolla, California) software. Data are presented as mean ± s.e.m. or median ± interquartile range as indicated. All data were tested for normality using the D’Agostino-Pearson omnibus normality test to guide the selection of parametric or non-parametric statistical tests. For parametric data, a two-tailed t test or two-way AN0VA was used, with a post hoc Sidak’s test for multiple comparisons. For non-parametric data, a two-tailed Mann-Whitney inverted U test was used. P<0.05 was considered significant.

## II. RESULTS

### i. Inhibitory synaptic transmission is impaired in adult Disc1df/+ mice

As a first estimation of inhibitory drive in the mPFC, we recorded mIPSCs onto L2/3 PNs in the dorsal anterior cingulate cortex (dACC) subregion of the mPFC in adult mice (postnatal day (P) 70). We found that, compared with wild type (WT) littermates, Disc1df/+ mice had significantly reduced mIPSC frequency (WT, 3.75 ± 3.25 Hz, n = 27 cells; Disc1df/+, 2.27 ± 2.72 Hz, n = 29 cells;U = 217.0, P<0.01, Mann-Whitney U test), but not amplitude (WT, 12.5 ± 2.48 pA, n = 29 cells; Disc1df/+, 12.52 ± l.5l pA, n = 27 cells; U = 351.0, P = 0.5l, Mann-Whitney U test) (Fig. 1A-D). The two groups did not differ in measures of miniature excitatory postsynaptic currents (mEPSC) (frequency: WT, 3.47 ± 1.97 Hz, n = 23 cells; Disc1df/+, 2.79 ± 2.4 Hz, n = 20 cells; U = 173.0, P = 0.17, Mann-Whitney U test; amplitude: WT, 8.91 ± 0.18 pA; Disc1df/+, 8.99 ± 0.37 pA, t(41) = 0.18, P = 0.86, t-test) (Fig. 1E-H).

Notably, we found that the frequency (albeit not amplitude) of mIPSCs recorded from dACC L2/3 PNs in Disc1df/+ mice was lower than their WT littermates at preweanling (~P15) age (Fig. 2). These data indicate that the inhibitory synaptic transmission is selectively impaired in the mPFC of Disc1df/+ mice, and that this impairment starts early in postnatal development.

### ii. Altered presynaptic function of PV interneurons in Disc1df/+ mice

A reduction in mIPSC frequency could result from a decrease in synaptic transmission from one or more inhibitory IN populations. To investigate the source of reduced inhibitory drive onto L2/3 PNs in the dACC of Disc1df/+ mice, we sought to examine the IPSCS originating from either PV or S0M INs. To this end, we selectively expressed channelrhodopsin (ChR2), the light-gated cation channel (Zhang et al., 2006), in PV or S0M INS by injecting the dACC of Disc1df/+; PV-Cre or Disc1df/+ S0M-Cre mice, as well as their Disc1+/+ (WT) littermates, with an adeno-associated virus (AAV) expressing ChR2 in a Cre-dependent manner (AAV-DIO-ChR2(H134R)-YFP). After viral expression had reached optimal levels, we prepared acute brain slices from these mice and recorded from dACC L2/3 PNs light-evoked IPSCs (Fig. 3A, D. We used paired light pulses (pulse duration1 ms) with an inter-pulse-interval of 50, 100, or 150 ms, and measured the ratio of the peak amplitude of the second IPSC over that of the first (IPSC2/IPSC1), also knownas paired-pulse ratio (PPR)(Fig. 3B, E). A similar technique has previously been used to interrogate presynaptic GABA release from PV interneurons (Chu, et al. 2012).

**Fig. 1.**
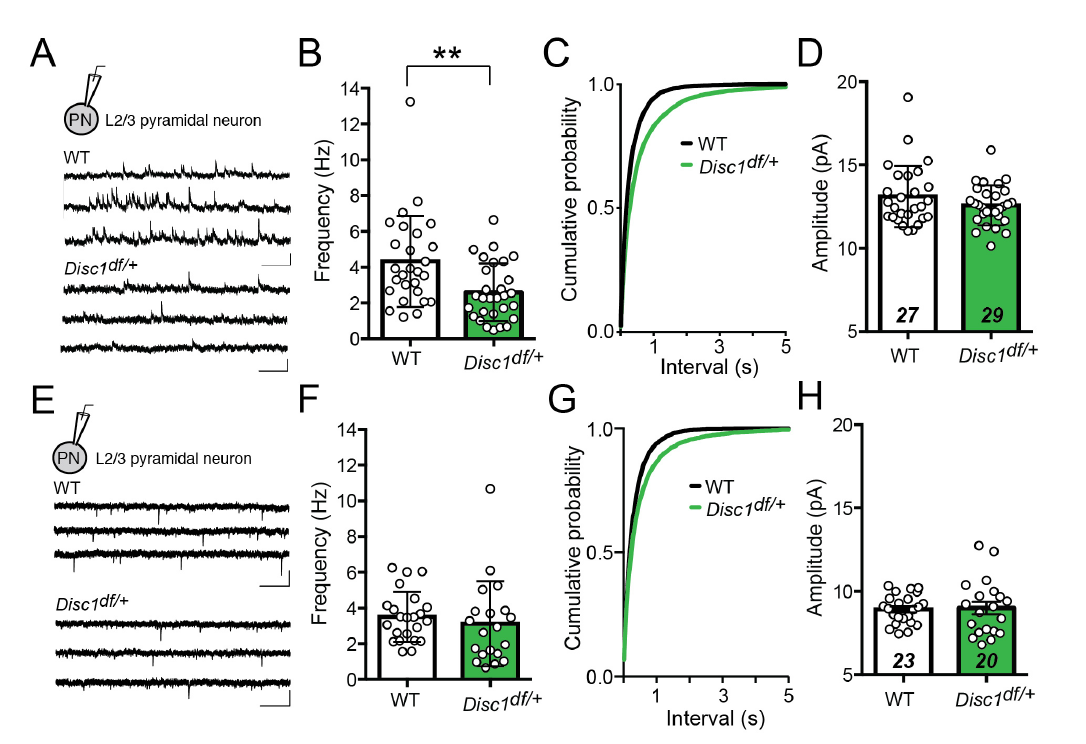
Reduced inhibitory synaptic transmission onto L2/3 pyramidal neurons in the mPFC of adult Disc1df/+ mice. A. Recording configuration and sample mIPSC traces recorded from L2/3 PNs in the mPFC of WT (upper) and Disc1df/+ (lower) mice at ~P70. B. Median mIPSC frequency (WT,n = 27 cells; Disc1df/+, n = 29 cells). C. Cumulative probability distributions of mIPSC inter-event-intervals. D. Median mIPSC amplitude (WT, n = 27 cells; Disc1df/+, n = 29 cells). E. Recording configuration and sample mEPSC traces recorded from L2/3 PNs in the mPFC of WT (upper) and Disc1df/+ (lower) mice. F. Median mEPSC frequency (WT, n = 23 cells; Disc1df/+, n = 20 cells). G. Cumulative probability distributions of mEPSC inter-event-intervals. H. Mean mEPSC amplitue (WT, n = 23 cells; Disc1df/+, n = 20 cells). All scale bars represent 20 pA, 500 ms. Bar graphs indicate median ± interquartile range (B,D,F) or mean ± s.e.m. (H), as appropriate. **P<0.01, Mann-Whitney U test.

We found that the PPR of GABAergic transmission between PV INs and L2/3 PNs was significantly increased in the Disc1df/+ mice compared with their WT littermates at the 50 and 100 ms inter-pulse-intervals (WT, n = 13; Disc1df/+, n = 10; interval: F(2, 42) = 6.77, P < 0.01; genotype: F(1, 21) = 10.77, P < 0.01; interaction: F(2, 42) = 3.92, P < 0.05; **P < 0.001; *P< 0.05; two-way repeated-measures (RM) ANOVA followed by Sidak’s tests) (Fig. 3C), suggesting that GABA release from PV INs is impaired. In contrast, the PPR of GABAergic synaptic transmission from SOM INs to L2/3 PNs did not differ between genotypes (WT, n = 15, Disc1df/+, n = 12; interval: F(2, 50) = 24.88, P <0.0001; genotype: F(1, 25) = 1.64, P = 0.21; interaction F(2, 50) = 0.47, P = 0.63, two-way RM ANOVA) (Fig. 3F). SOM-evoked IPSCs displayed significantly slower decay kinetics than PV-evoked IPSCs (Fig. 3G, H), consistent with previous reports (Koyanagi et al., 2010; Ma et al., 2012). No differences in IPSC kinetics were observed between Disc1df/+ mice and their WT littermates (cell type: F(1, 46) = 90.82, ****P<0.0001; genotype: F(1,46) = 0.678, P = 0.41; two-way ANOVA) (Fig. 3G, H). In light of the observed reduction in mIPSC frequency, the increased PPR of PV-mediated IPSCs suggests that there is a presynaptic deficit in GABA release from PV cells to L2/3 PNs in the mPFC of Disc1df/+ mice.

**Fig. 2.**
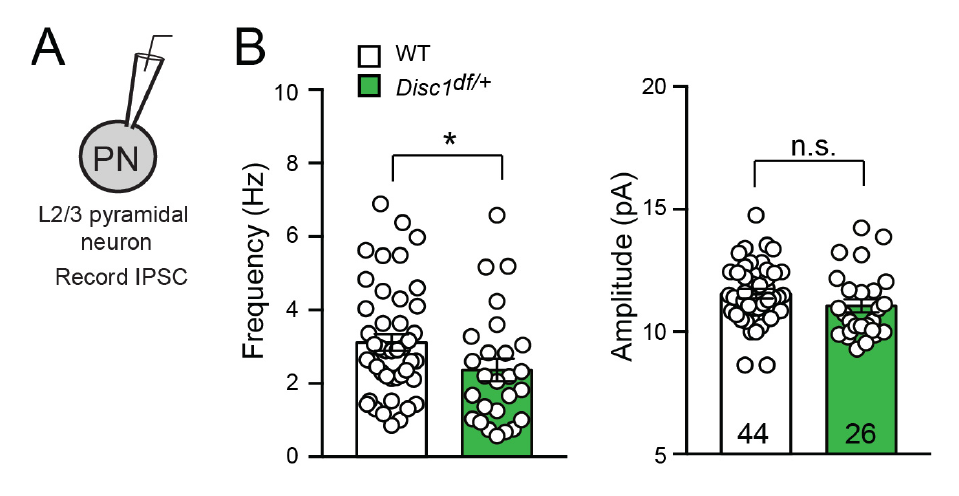
Reduced inhibitory synaptic transmission onto L2/3 pyramidal neurons in the mPFC of preweanling (~P15) Disc1df/+ mice. A. Recording configuration of mIPSC from L2/3 PNs in the mPFC. B. Quantification of mIPSC frequency (left) (WT, n = 44 cells; Disc1df/+, n = 26 cells;*P<0.05, Mann-Whitney U test) and amplitude (right) (WT, n =; 44 cells; Disc1df/+, n = 26 cells;.P = 0.12, t test). Frequency is presented as median ± interquartile range and amplitude is presented as mean ± s.e.m.

### iii. Reduced feedforward inhibition in a thalamus-mPFC circuit in Disc1df/+ mice

The mediodorsal nucleus of the thalamus (MD) sends major projections to the mPFC. This MD-mPFC circuit has been implicated in cognitive processes such as working memory and executive function that are impaired in schizophrenia (Lesh et al., 2011; Parnaudeau et al., 2013; Parnaudeau et al., 2015). We recently reported that the excitatory inputs from the MD drive mPFC feedforward inhibition(FFI)that is selectively mediated by mPFC PV INs (Delevich et al., 2015). Given the deficit in GABA release from PV INs to PNs in the mPFC of Disc1df/+ mice (Fig. 3), we reasoned that in these mice the FFI in the MD-mPFC circuit is affected. To test this hypothesis,we injected the MD of Disc1df/+ mice and their WT littermates with AAV-ChR2(H134R)-YFP. After viral expression reached optimal levels we used these mice to prepare acute brain slices, in which we recorded both excitatory and inhibitory synaptic transmission onto dACC L3 PNs in response to optogenetic stimulation of MD axons (Fig. 4A-F).

Brief (0.5 ms) light stimulation evoked monosynaptic EPSCs and disynaptic IPSCs in L3 PNs in the dACC (Fig. 4C) (and see (Delevich et al., 2015)). The latencies and kinetics of the EPSCs and IPSCs in Disc1df/+ mice were similar to those in WT mice (Fig. 5). However, we found that the contribution of inhibitory synaptic transmission to total synaptic inputs, measured as IPSC-peak/(IPSCpeak+EPSCpeak), or Ipeak/(Ipeak+Epeak), was significantly lower in Disc1df/+ mice than in WT mice when comparing the means of the two groups of animals (Disc1df/+, 0.52 ± 0.03, n = 11 mice; WT, 0.70 ± 0.02, n = 14 mice; t(23) = 5.73, ****P<0.0001, t test) (Fig. 4D), or the means of the two groups of neurons (Disc1df/+, 0.60 ± 0.03; n = 30 cells, WT, 0.70 ± 0.02, n = 40 cells; t(68) = 3.17, **P<0.01, t test) (Fig. 4E). In addition, the slope of a linear regression describing the relationship between IPSCs and EPSCs of individual neurons in the Disc1df/+ mice was lower than that in the WT (Disc1df/+, = 30 neurons, WT, = 40 neurons) (Fig. 4F). These results together indicate that the MD-driven FFI in the mPFC is reduced by DISC1 deficiency.

**Fig. 3.**
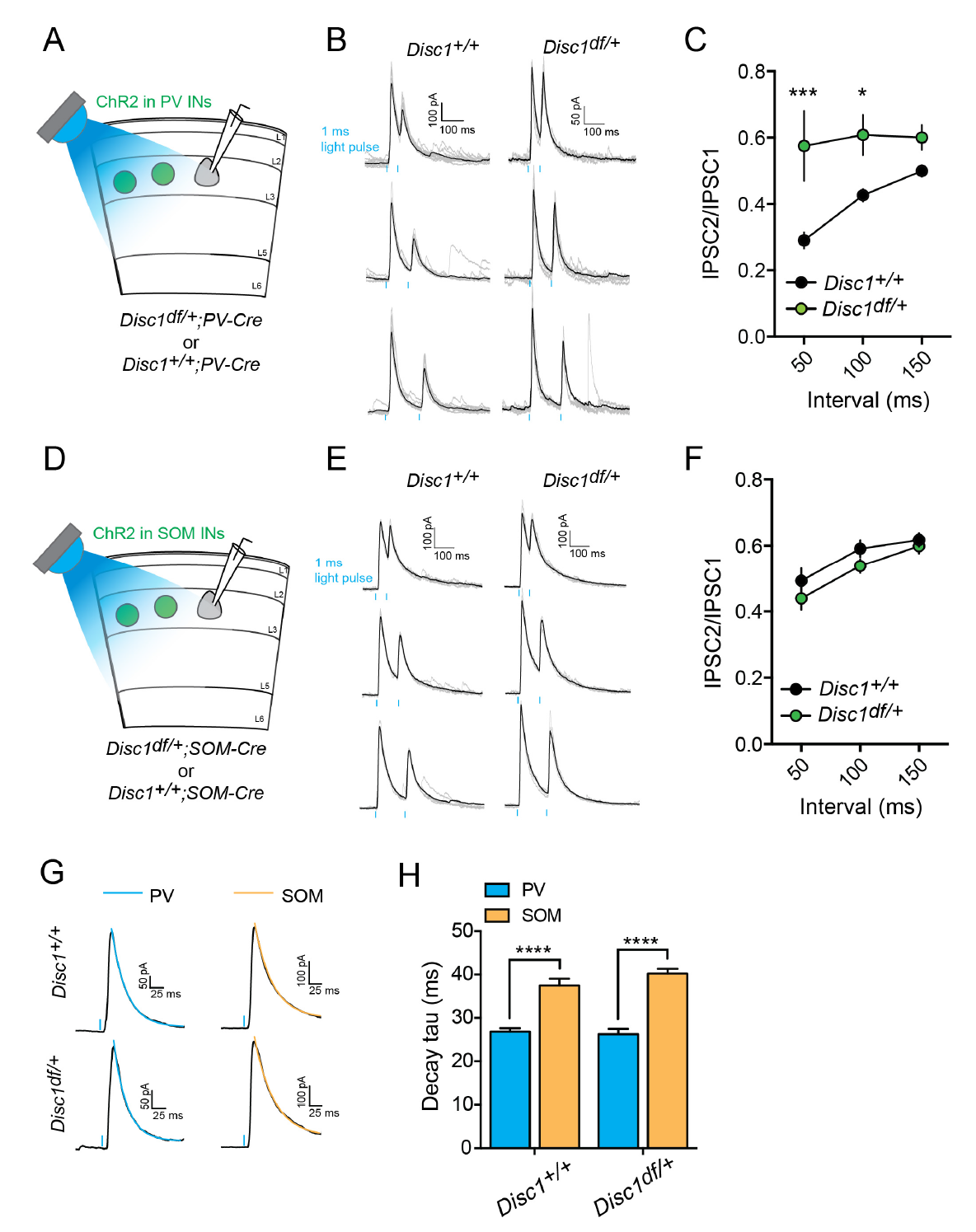
Impaired presynaptic function of PV but not SOM INs in the mPFC of Disc1df/+ mice. A. Schematic of the experimental configuration. B. Sample traces of PV IN-mediated IPSCs recorded from WT (left panel) or Disc1df/+ (right panel) mice. Paired light pulses (1 ms duration; blue bars) was delivered at an interval of 50 ms (top), 100 ms (middle) or 150 ms (bottom). C. Quantification of PPR for each genotype (WT, n = 13 cells; Disc1df/+, n = 10 cells). *P<0.05, ***P<0.001, two-way ANOVA followed by Sidak's test. D. Schematic of the experimental configuration. E. Sample traces of SOM IN-mediated IPSCs recorded from WT (left panel) or Disc1df/+ (right panel) mice. Paired light pulses (1 ms duration; blue bars) was delivered at an interval of 50 ms (top), 100 ms (middle) or 150 ms (bottom). F. Quantification of PPR for each genotype (WT, n = 15 cells, Disc1df/+, n = 12 cells). G. Sample IPSC traces evoked by optogenetic stimulation of PV or SOM INs. Colored lines indicate exponential fits to the decays of the IPSCs. H. Quantification of IPSC decay tau. ****P<0.0001, t test. Data in C, F, and H are presented as mean ± s.e.m.

### iv. Enhanced input but reduced output of PV interneurons in Disc1df/+ mice

The decrease in FFI in the MD-mPFC pathway in Disc1df/+ mice could result from the impairment in GABA release from PV INs in the mPFC (Fig. 3), or other potential deficits in these mice, such as a reduction in MD recruitment of mPFC PV INs. To disambiguate these possibilities, we first examined MD recruitment of mPFC PV INs and PNs. We recorded EPSCs from PV IN and PN pairs in the dACC in response to optogenetic stimulation of MD axons (Fig. 6A). We found that in WT mice, the amplitudes of these thalamocortical EP-SCs were similar between PV INs and adjacent PNs (PV,−109.3 ± 109.7 pA, PN, −129.1 ± 103.2 pA, n = 15 pairs, W = 0, p = 1.0, Wilcoxon matched-pairs signed rank test) (Fig. 6B, C). By contrast, in Disc1df/+ mice, the thalamocortical EPSCS onto PV INS were much larger than those onto neighboring PNs (PV, −153.4 ± 211.9 pA; FN, −73.74 ± 104.6 pA; n = 14 pairs, W = −83, **p < 0.0l Wilcoxon matched-pairs signed rank test) (Fig 6B, C). Thus, the MD-driven excitatory drive onto PV INS is enhanced relative to that onto PNs in the mPFC of Disc1df/+ mice. This change cannot account for the decrease in FFI in the MD-mPFC circuit in these mice (Fig. 4).

**Fig. 4.**
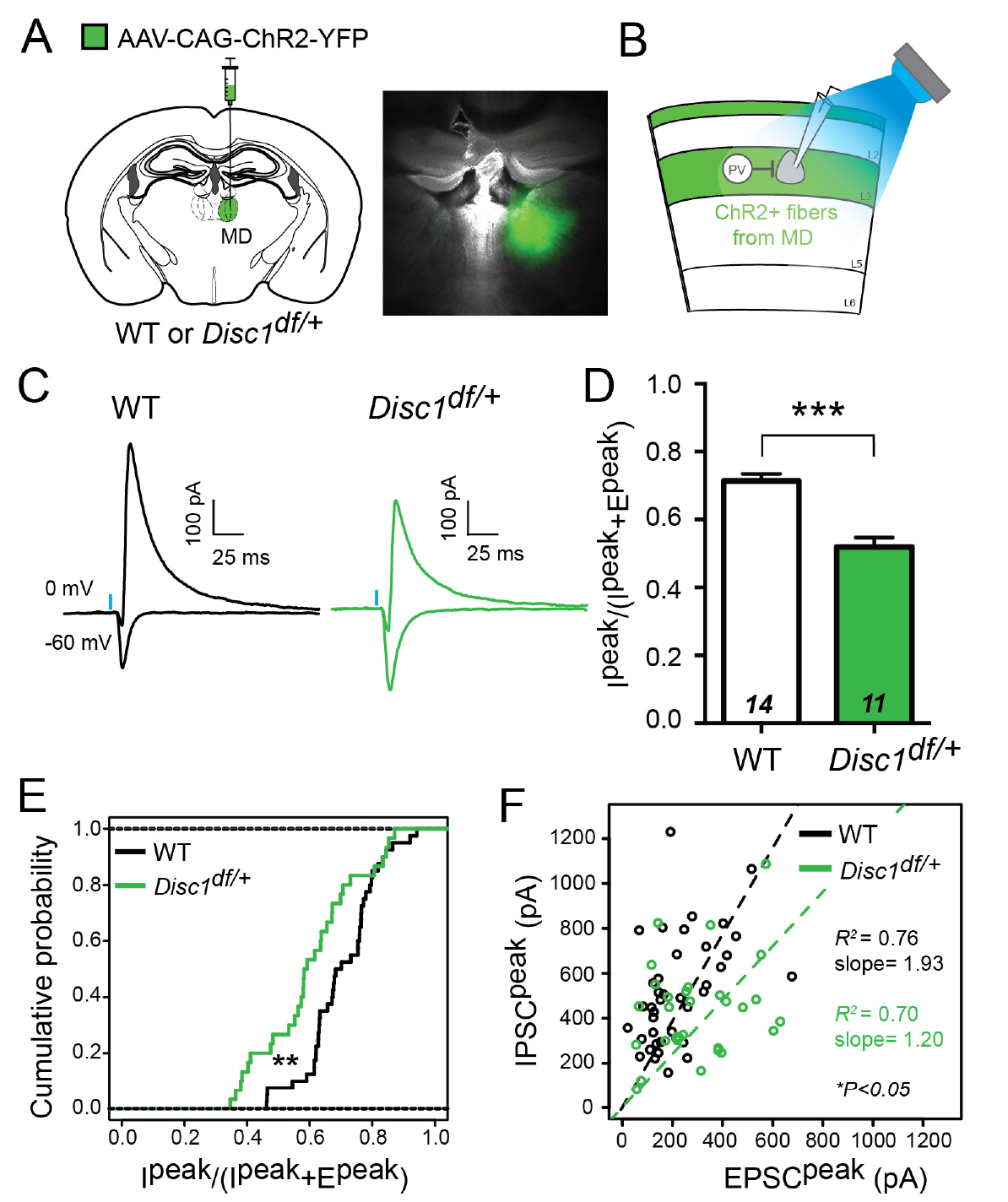
Reduced feedforward inhibition in the MD-mPFC circuit in Disc1df/+ mice. A. & B. Schematics of the experimental configuration. The right panel of (A) is an image of a brain section from a mouse used in electrophysiological recording, showing the MD infected with AAV-CAG-ChR2-YFP. C. Representative traces of EPSC (recorded at −60 mV) and IPSC (recorded at 0 mV) from L3 PNs. D. To estimate the relative recruitment of disynaptic FFI versus monosynaptic excitation, we divided peak IPSC (I^peak^) by the sum of peak IPSC and peak EPSC (I^peak^ + E^peak^). WT, n = 14 mice, Disc1df/+, n = 11 mice; ****P < 0.001, t test. E. Same as in (D), except that the cumulative probability distributions of the values for individual neurons are shown. WT, n = 40 cells, Disc1df/+, n = 30 cells; **P < 0.01, Kolmogorov-Smirnov test. F. Scatter plot showing the peak amplitudes of IPSC and EPSC for individual neurons. Each circle represents one neuron (WT, n = 30 cells; Disc1df/+, n = 40 cells). Dashed lines are linear regression lines for neurons in WT mice and Disc1df/+ mice. The slopes of the regression lines significantly differed at the 0.95 confidence level (*P < 0.05). Data in (D) are presented as mean ± s.e.m.

Notably, we found that this FFI PPR was significantly higher in Disc1df/+ mice than in the WT (WT, 0.0 ± 0.1 (median ± interquartile range), n = 24 cells; Disc1df/+, 0.24 ± 0.34, n = 17 cells; U = 62, ****p = 0.0001, twotailed Mann Whitney U test) (Fig. 6D, E). Note to reduce variability in measuring the PPR, we set the light-stimulation such that there was no difference between genotypes in the average amplitude of the first IPSCS (WT, 439.3 ± 41.31 pA, n = 24 cells; Disc1df/+, 367.1 ± 46.26 pA, n = 17 cells; t(39) = 1.152, P = 0.256, unpaired t test) (Fig. 6F). These results lend further support to the finding that presynaptic function of PV INS is altered in Disc1df/+ mice (Fig. 3). Together, our data suggest that in Disc1df/+ mice, GABA release from prefrontal PV INS is reduced, leading to decreased FFI in the MD-mPFC circuit.

## III. DISCUSSION

It has been hypothesized that the cognitive deficits of schizophrenia may be the consequence of imbalanced excitation and inhibition (E-I) in key neural circuits (Kehrer et al., 2008; Lisman, 2012; Marin, 2012). Consistent with this hypothesis, several studies have shown that imposing elevated E/I ratio within the prefrontal cortex impairs cognitive processing in rodents (Cho et al., 2015; Yizhar et al., 2011). This E-I imbalance can arise from either an increase in excitation or a reduction in inhibition. Here, we examined synaptic transmission in the MD-mPFC circuit in mice deficient for the Disc1 gene, the human homolog of which has been shown to be a rare genetic risk factor for major psychiatric disorders, including schizophrenia (Millar et al., 2000). We found that DISC1deficiency caused E-I imbalance, measured as decreased FFI, in the MD-mPFC circuit. Furthermore, this effect could be accounted for by a reduction in GABA release from PV INs in the mPFC.

**Fig. 5.**
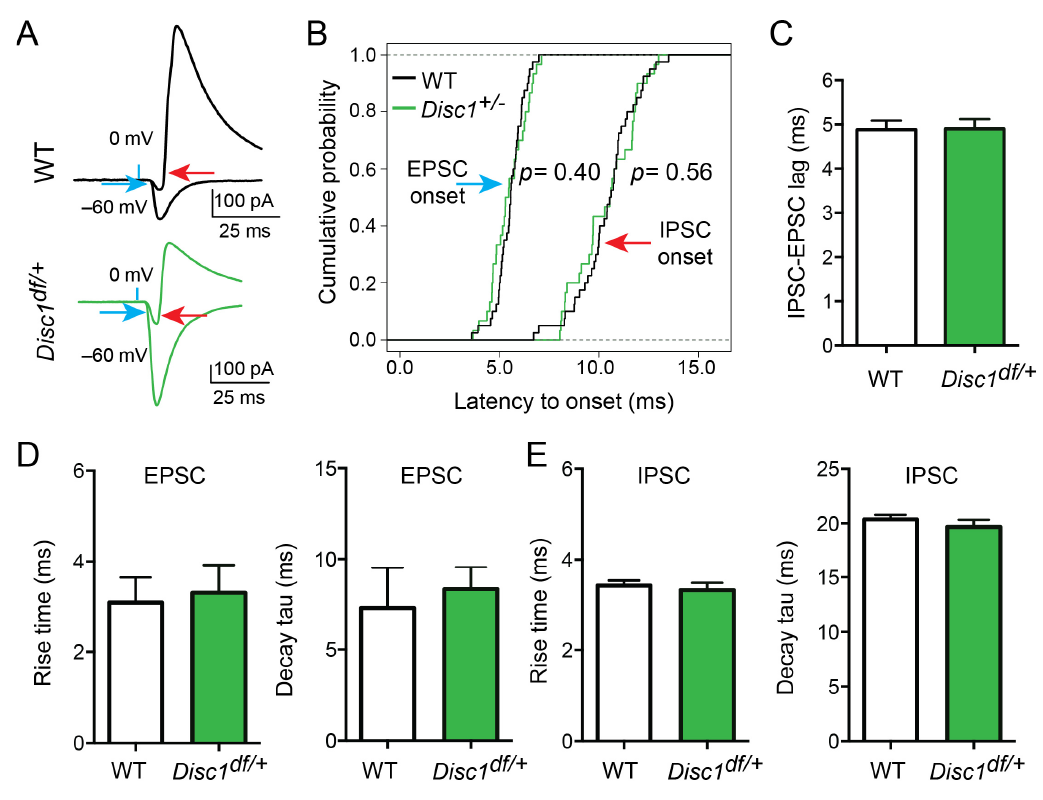
Normal latencies and kinetics of synaptic transmission onto L2/3 pyramidal neurons in the mPFC of Disc1df/+ mice. A. Sample traces of IPSC (recorded at 0 mV) and EPSC (recorded at −60 mV) recorded from L2/3 PNs in response to light-stimulation (blue bars) of MD axons. The latency to onset was measured from the time the light stimulus was triggered to the 10% EPSC (blue arrow) or IPSC (red arrow) rise time. Note that IPSC rise time was calculated from the peak of the inward current recorded at 0 mV. B. Cumulative probability distributions for EPSC latency to onset (left) and IPSC latency to onset (right) (EPSC, WT, n = 40 cells, Disc1df/+, n = 30 cells, P = 0.40; IPSC, WT, n = 40 cells, Disc1df/+, n = 30 cells, P = 0.56; Kolmogorov-Smirnov test). C. Quantification of IPSC-EPSC lag, calculated as the difference in the latency to onset between the IPSC and the EPSC of each neuron (see also A.) (WT, n = 40 cells, Disc1df/+, n = 30 cells; P > 0.05, t test). D. Quantification of the 10–90% EPSC rise time (left) and decay tau (right) (WT, n = 40 cells, Disc1df/+, n = 30 cells; P > 0.05, Mann-Whitney U test). E. Quantification of the 10–90% IPSC rise time (left) and decay tau (right) (WT, n = 40 cells, Disc1df/+, n = 28 cells; P > 0.05, t test). Data are presented as median ± interquartile range (D) or mean ± s.e.m. (C, E).

Interestingly, we observed that excitatory synaptic transmission onto PV cells driven by MD inputs was actually enhanced in Disc1df/+ mice. Therefore, there might be compensatory upregulation of excitatory inputs onto PV INs in response to the decrease in output from these neurons. A recent study investigating the same Disc1df/+ model reported that there is no change in the number of PV INs in the mPFC (Seshadri et al., 2015). Together, these findings suggest that the PV⤑PN synapses are the primary site of impairment in the MD-mPFC circuit in the Disc1df/+ mice.

**Fig. 6.**
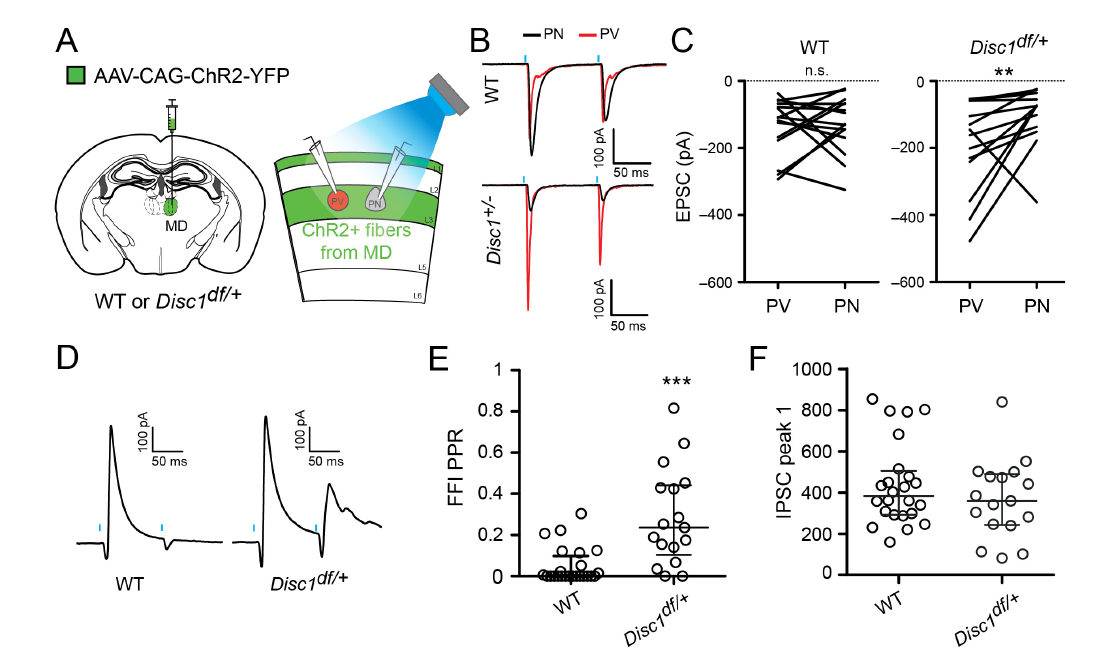
Impaired presynaptic function of PV INs underlies the deficit of FFI in Disc1df/+ mice. A. Left: a schematic of the experimental configuration. Right: a schematic of the recording configuration in the mPFC acute slices. A Tdtomato+ PV IN (red) and an adjacent PN (gray) in L3 of the mPFC were recorded simultaneously or sequentially. EPSCs onto these neurons were evoked by optogenetic stimulation (0.5 ms light pulses; blue bars) of MD axons. B. Sample EPSC traces recorded from PV IN and PN pairs are superimposed and color-coded. C. Quantification of the EPSC peak amplitude. n.s., not significant (P = 1.0); **P < 0.01; Wilcoxon matched-pairs signed ranks test. D. Sample traces of FFI currents recorded from L3 PNs in response to optogenetic stimulation of MD axons. E. Quantification of PPR of the MD-driven FFI onto L3 PNs. ***P < 0.001, Mann-Whitney U test. F. The mean amplitude of the first IPSC is consistent between genotypes. Data in E and F are presented as median ± interquartile range.

While our study is the first to specifically detect a presynaptic deficit in PV INs in a DISC1 genetic deficiency model, previous studies using different DISC1 models have reported that DISC1 influences inhibitory IN function or development: spontaneous IPSC frequency is reduced in the frontal cortex of male mice expressing a truncated mouse DISC1 (Holley et al., 2013); PV IN function is impaired in the mPFC of mice overexpressing a truncated DISC1 (Sauer et al., 2015); PV expression is reduced in the PFC of several DISC1 mouse models (Ayhan et al., 2011; Hikida et al., 2007; Lee et al., 2013; Niwa et al., 2010; Shen et al., 2008); and tangential migration of MGE-derived neurons is impaired by Disc1 mutation or RNA interference (Lee et al., 2013; Steinecke et al., 2012). These findings provide converging evidence that DISC1 perturbation affects prefrontal cortical inhibition, in particular that mediated by PV INs.

Abnormal functional connectivity between the MD and the prefrontal cortex (PFC) has been reported in patients with schizophrenia (Minzenberg et al., 2009; Mitelman et al., 2005; Seidman et al., 1994; Welsh etal., 2010; Woodward et al., 2012; Zhou et al., 2007) and has been suggested to be a potential biomarker of the disease (Anticevic et al., 2014). The coordinated activity between the MD and the PFC is important for working memory and flexible goal-oriented behavior (Mitchell and Chakraborty, 2013; Parnaudeau et al., 2013; Parnaudeau et al., 2015), faculties that are impaired in schizophrenia. 0ur findings suggest that impaired prefrontal PV IN function might contribute to the changes in MD-PFC communication in subjects with schizophrenia. These results support an emerging hypothesis from the human literature that local disinhibition of PFC may destabilize the flow of information through the thalam-ofrontal loop in schizophrenia (Anticevic et al., 2012). Given that few treatment options exist to address the cognitive symptoms of schizophrenia and related disorders, efforts towards understanding the cellular and molecular mechanisms underlying abnormal thalam-ofrontal functional connectivity may yield therapies that will improve patient outcomes.

